# libsbmljs — Enabling Web–Based SBML Tools

**DOI:** 10.1101/594804

**Authors:** J Kyle Medley, Joseph Hellerstein, Herbert M Sauro

## Abstract

The SBML standard is used in a number of online repositories for storing systems biology models, yet there is currently no Web–capable JavaScript library that can read and write the SBML format. This is a severe limitation since the Web has become a universal means of software distribution, and the graphical capabilities of modern web browsers offer a powerful means for building rich, interactive applications. Also, there is a growing developer population specialized in web technologies that is poised to take advantage of the universality of the web to build the next generation of tools in systems biology and other fields. However, current solutions require server– side processing in order to support existing standards in modeling. We present libsbmljs, a JavaScript / WebAssembly library for Node.js and the Web with full support for all SBML extensions. Our library is an enabling technology for online SBML editors, model–building tools, and web–based simulators, and runs entirely in the browser without the need for any dedicated server resources. We provide NPM packages, an extensive set of examples, JavaScript API documentation, and an online demo that allows users to read and validate the SBML content of any model in the BioModels and BiGG databases. We also provide instructions and scripts to allow users to build a copy of libsbmljs against any libSBML version. Although our library supports all existing SBML extensions, we cover how to add additional extensions to the wrapper, should any arise in the future. To demonstrate the utility of this implementation, we also provide a demo at https://libsbmljsdemo.github.io/ with a proof–of–concept SBML simulator that supports ODE and stochastic simulations for SBML core models. Our project is hosted at https://libsbmljs.github.io/, which contains links to examples, API documentation, and all source code files and build scripts used to create libsbmljs. Our source code is licensed under the Apache 2.0 open source license.

## Introduction

The SBML (1) standard is used for encoding reaction network models in systems biology research in a reusable, exchangeable, and future–proof manner. One of the factors behind SBML’s wide adoption is the SBML standard’s process for introducing extension modules, which allow incremental incorporation of new capabilities. While the core components of the standard are designed for describing kinetic chemical reaction network models, SBML extensions exist for encoding constraint–based models (the “flux– balance constraints” extension, employed by the widely used COBRA framework for constraint–based modeling (2, 3)), and rule–based models (the SBML “multi” extension (4)). SBML is used in several online model repositories including BioModels (5, 6) and JWS Online (7, 8), which host primarily kinetic reaction network models, and BiGG Models cite (9), which hosts primarily genome–scale constraint– based models.

Despite this wide–spread adoption and inclusion in several online repositories, no feature–complete JavaScript library currently exists that can run in a web browser (a native Node.js module exists, but cannot run in the browser). Thus, these online repositories must rely on server–side processing of all SBML–related requests. A JavaScript library would allow these services to offload some of their processing to the client, and would also allow for more interactive features on the Web. Furthermore, the Web is becoming a major platform for systems biology tools. With the advent of Web applications for pathway visualization (Escher (10)), gene interaction network visualization (Cytoscape.js (11)), expression analysis (ZBIT (12)) and integrated design systems (Caffeine (13)), the need for a JavaScript library which can read and write SBML becomes imperative.

We present libsbmljs, a feature–complete JavaScript library for reading and writing SBML in the browser and Node.js. libsbmljs uses the full codebase of the libSBML C++ library compiled to the web using Emscripten, a toolset for compiling C++ projects to the web. Emscripten emits WebAssembly (14), a W3C standard for running platform– independent binary code on the web that is supported on all major browsers. We have designed a JavaScript wrapper around this binary format that allows libsbmljs to be used like a normal JavaScript library. Our wrapper supports *all* SBML Level 3 extensions, meaning it can read and write any type of SBML content. Since our library runs in the browser, it does not require a dedicated web server. This is an important consideration for academic software, where long–term maintenance cost is a concern.

## Methods

Prior work on implementing the SBML standard has resulted in two libraries: libSBML (15), a C++ library with interfaces for many languages, and JSBML (16, 17), a platform– independent pure Java library. While the existence of these separate implementations is certainly a convenience for C++ and Java developers respectively, it necessitates the maintenance of two independent libraries. Rather than attempt to create a third implementation in pure JavaScript, we have created a web–capable interface for the libSBML C++ library using Emscripten (18), a C++–to–JavaScript compiler. Despite its C++ origins, libsbmljs is completely platform independent and runs on modern browsers on any device which supports web standards.

Compiling a C++ library with Emscripten does not produce a ready–to–use JavaScript library automatically. Instead, Emscripten compiles to WebAssembly (19), a low–level binary format similar to x86 machine code but with additional features for security and platform–independence. Since WebAssembly is very low level, it is difficult to use to design JavaScript web applications. Instead, Emscripten can be used to also compile a JavaScript interface that abstracts the low– level details of calling into WebAssembly and instead allows developers to use familiar JavaScript objects and methods. However, this interface is not generated automatically by Emscripten. Instead, it must be manually specified using WebIDL.

Web IDL is a World Wide Web Consortium (W3C^®^) standard that specifies *interfaces* to EMCAScript (i.e. JavaScript) objects. For example, the libSBML C++ class `SBase` has the method `getId()`, which returns a string. In WebIDL, this would be specified as:

~~~
/**
 * SBase: the base class of
 * most SBML elements
 */
[**Prefix**=**“libsbml::”**]
**interface SBase** {
 /**
  * Returns this element’s
  * id attribute
  */
 **DOMString getId**();
};
~~~

In the example above, the body of the `getId` method is intentionally left blank because it will delegate to the corresponding WebAssembly routine. Using syntax similar to the above, we manually designed WebIDL interface files for every libSBML class and method. However, one issue remains with this approach. The comments entered into the IDL definition above will not appear in the JavaScript interface generated by Emscripten. Thus, there is no way of adding documentation to the generated JavaScript code, which defeats any attempt to generate API documentation. To remedy this issue, we created a script to automatically extract documentation strings from IDL files and insert them into the generated JavaScript code. This allowed us to generate extensive API documentation using `documentationjs`, a documentation generator for JavaScript.

### Special Considerations for Usage

Emscripten– generated WebAssembly/JavaScript libraries are supported on a wide variety of browsers and devices (https://github.com/libsbmljs/libsbmljs lists the browsers we have tested). However, there are minor differences between these libraries and regular JavaScript libraries, which are described below.

#### Asynchronous Loading

Emscripten–generated libraries load asynchronously. In other words, the library cannot be used immediately as soon as the web page has loaded. This is due to the fact that Emscripten–generated libraries consist of both a JavaScript source file (.js) containing JavaScript classes and methods, and a WebAssembly file (.wasm) containing the compiled C++ code. The browser may load the JavaScript source file before completely loading and compiling the WebAssembly file. In order to accommodate this, Emscripten libraries provide a ‘then()’ method for the JavaScript module object similar to a JavaScript Promise. This method accepts a callback that will execute once the WebAssembly is fully downloaded and compiled.

#### Manual Memory Management

Most modern languages feature some type of automatic garbage collection. However, WebAssembly is a low–level binary–like format, and hence does not provide high–level features like garbage collection. This means that whenever the user creates an object in libsbmljs using the new keyword, the user must also destroy the object using `libsbml.destroy(obj)`. In most cases, this simply amounts to destroying the SBML document when it is no longer needed.

In terms of modern programming languages, this may seem like a significant regression, but it is an unavoidable tradeoff when using C++ compiled WebAssembly, at least for currently available technology (a proposal exists to add garbage collection to WebAssembly (20), but an implementation is not available at the time of writing). In the event that the user forgets to call the `libsbml.destroy` function, the allocated object will persist in the browser’s memory until the browser tab is closed. Since our main target users are developers of web applications, and browser tabs are short–lived, we do not believe this is a significant concern. However, Node.js developers should take care to destroy all created objects. The same requirement also applies to libSBML’s native Node.js module.

#### Client–side SBML Simulation

We used libsbmljs to construct a fully client–side, web–based SBML simulator `(sbml_websim)`, which supports ODE–based (using the Bulirsch–Stoer algorithm (22, 23) implemented in JavaScript (24)) and stochastic (using the Next Reaction / “Gibson” method (25)) simulations. This simulator is directly connected to the BioModels / BiGG Models browser demo associated with this manuscript (https://libsbmljsdemo.github.io). The simulator supports SBML core features including rate rules and events. Although slow compared to state–of–the–art simulators like libroadrunner (26), the main utility of this simulator is to demonstrate the technological readiness of this approach for creating standards–supporting web–apps. This method could be used, for example, to integrate simulation capability into the BioModels and BiGG Models repositories. Figure 2 shows web–based simulations of each of these models using `sbml_websim`. Figure 3 shows an example stochastic simulation.

**Fig. 1.**
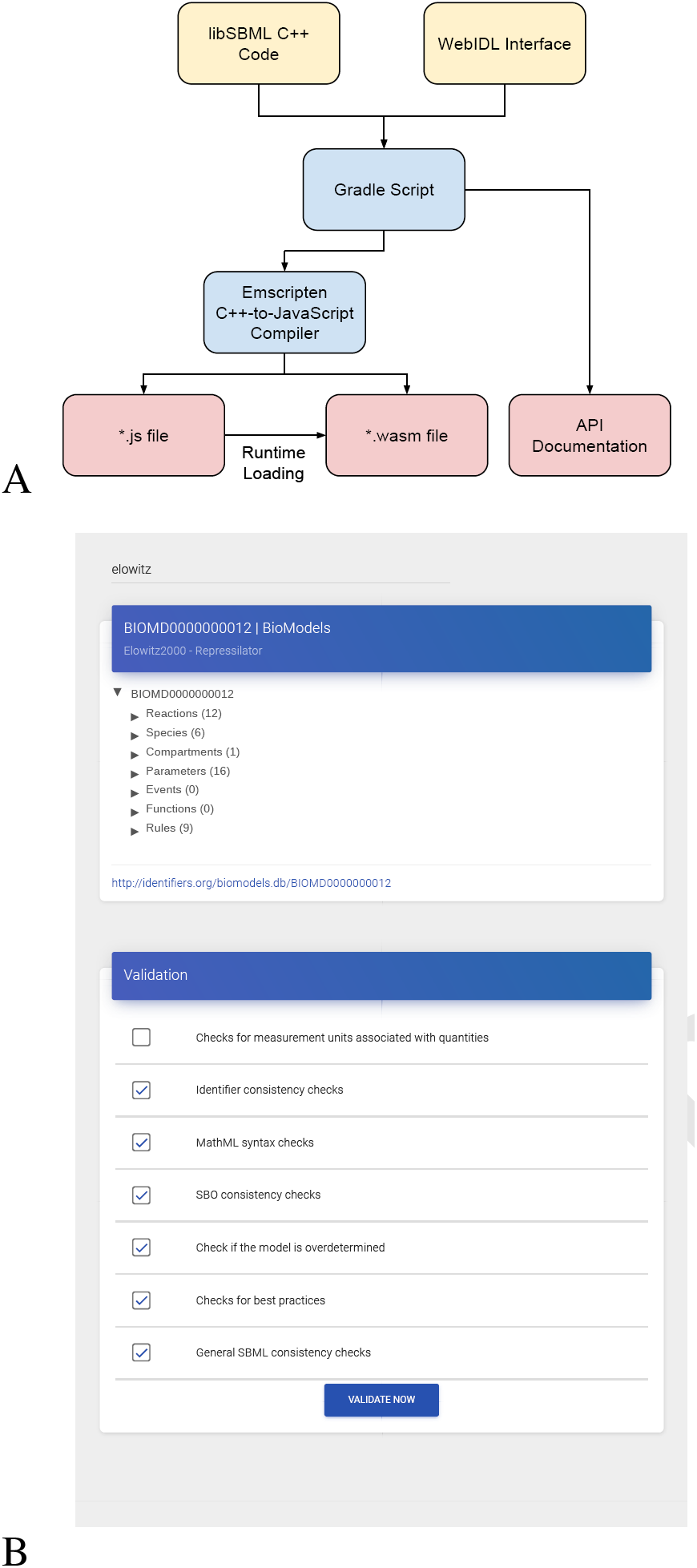
**(A)** A workflow diagram of the process used to produce libsbmljs. The lib-SBML C++ source code and a hand–written WebIDL interface are processed by a Gradle script to produce Emscripten–compiled bytecode and JavaScript API documentation. The Emscripten bytecode is further compiled into separate JavaScript (*.js) and WebAssembly (*.wasm) files. When the JavaScript source file is loaded by the browser, it executes instructions to fetch the corresponding WebAssembly file asynchronously. These two files are then combined into an npm package. **(B)** A screenshot of the demo page showing the Repressilator model (21) in the BioModels database (BIOMD0000000012). After selecting a model via using the demo’s search bar or uploading an SBML file, the demo allows the user to view SBML content as a tree–like structure and validate the SBML model subject to the validation options provided by libSBML. This particular model can be viewed at https://libsbmljsdemo.github.io/#/view?m=BIOMD0000000012

**Fig. 2.**
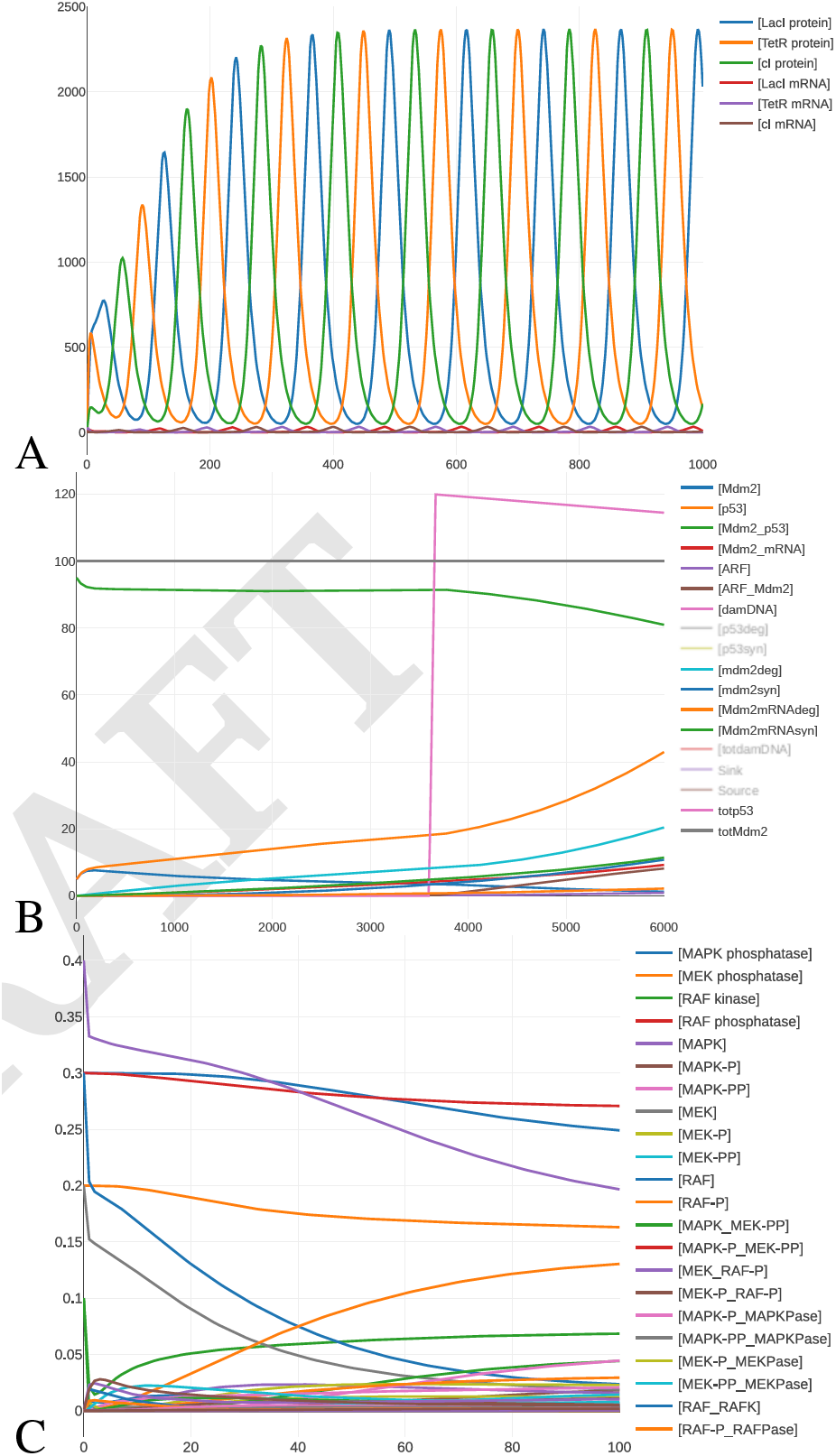
Example web–based simulations of BioModels: the repressilator (https://libsbmljsdemo.github.io/#/view?m=BIOMD0000000012, 12 reactions, **A**), p53 p14ARF (https://libsbmljsdemo.github.io/#/view?m=BIOMD0000000189, 14 reactions, **B**), and MAPK (https://libsbmljsdemo.github.io/#/view?m=BIOMD0000000014, 300 reactions, **C**) models. These results were separately compared to the libroadrunner simulator to verify accuracy (not shown).

**Fig. 3.**
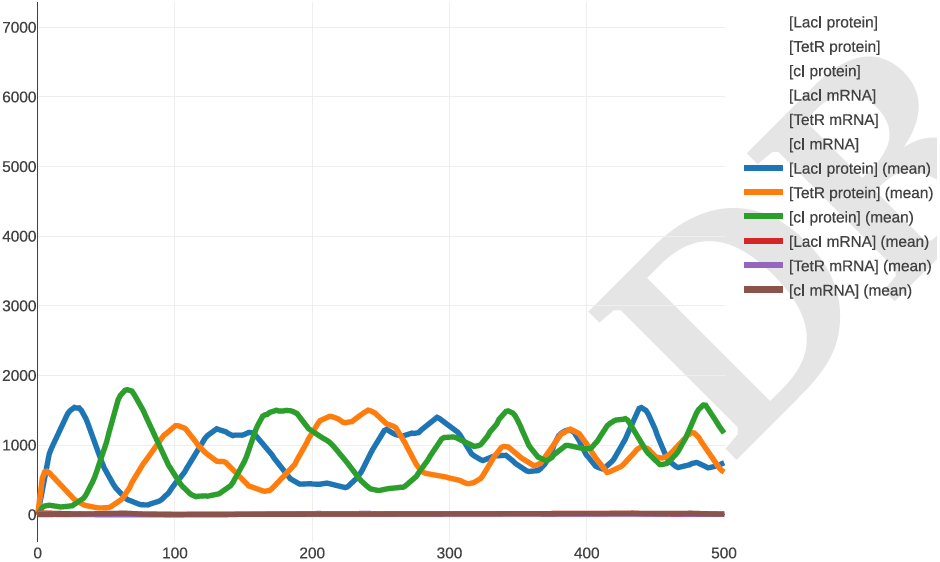
A stochastic simulation of the repressilator model the Next Reaction Method(25). `sbml_websim` allows the user to repeat the stochastic simulation for a desired number of replicates (10 here) and plots all replicates (faint lines) in addition to the mean value of each variable over all replicates (solid line).

## Discussion & Conclusion

Currently, there is no web–capable library that can read and write SBML models. We have presented a WebAssembly / JavaScript library that can read and write all SBML packages. We have provided tutorials, examples and extensive API documentation for potential users. We have also provided a modular build system that can be used to regenerate the wrapper from any recent checkout of the libSBML C++ library from the stable or experimental branch, as well as in– browser tests of the wrapper using the Karma testing engine. Additionally, we have used this wrapper to create the first web–based client–side SBML simulator. We hope these advances will enable the development of systems biology web applications and services that can use the SBML standard.

## ACKNOWLEDGEMENTS

We would like to thank Ricardo Henriques for providing the template on which this article is based.

## Bibliography

1. Michael Hucka, Andrew Finney, Herbert M Sauro, Hamid Bolouri, John C Doyle, Hiroaki Kitano, Adam P Arkin, Benjamin J Bornstein, Dennis Bray, Athel Cornish-Bowden, et al. The systems biology markup language (sbml): a medium for representation and exchange of biochemical network models. Bioinformatics, 19(4):524–531, 2003.

2. Scott A Becker, Adam M Feist, Monica L Mo, Gregory Hannum, Bernhard Ø Palsson, and Markus J Herrgard. Quantitative prediction of cellular metabolism with constraint-based models: the cobra toolbox. Nature protocols, 2(3):727, 2007.

3. Jan Schellenberger, Richard Que, Ronan MT Fleming, Ines Thiele, Jeffrey D Orth, Adam M Feist, Daniel C Zielinski, Aarash Bordbar, Nathan E Lewis, Sorena Rahmanian, et al. Quantitative prediction of cellular metabolism with constraint-based models: the cobra toolbox v2. 0. Nature protocols, 6(9):1290, 2011.

4. Fengkai Zhang and Martin Meier-Schellersheim. Sbml level 3 package: multistate, multicomponent and multicompartment species, version 1, release 1. Journal of integrative bioinformatics, 15(1), 2018.

5. Nicolas Le Novere, Benjamin Bornstein, Alexander Broicher, Melanie Courtot, Marco Donizelli, Harish Dharuri, Lu Li, Herbert Sauro, Maria Schilstra, Bruce Shapiro, et al. Biomodels database: a free, centralized database of curated, published, quantitative kinetic models of biochemical and cellular systems. Nucleic acids research, 34(suppl 1): D689–D691, 2006.

6. Chen Li, Marco Donizelli, Nicolas Rodriguez, Harish Dharuri, Lukas Endler, Vijayalakshmi Chelliah, Lu Li, Enuo He, Arnaud Henry, Melanie I Stefan, et al. Biomodels database: An enhanced, curated and annotated resource for published quantitative kinetic models. BMC systems biology, 4(1):92, 2010.

7. Brett G Olivier and Jacky L Snoep. Web-based kinetic modelling using jws online. Bioinformatics, 20(13):2143–2144, 2004.

8. Martin Peters, Johann J. Eicher, David D. van Niekerk, Dagmar Waltemath, and Jacky L. Snoep. The jws online simulation database. Bioinformatics, 33(10):1589–1590, 2017. doi: 10.1093/bioinformatics/btw831.

9. Zachary A King, Justin Lu, Andreas Dräger, Philip Miller, Stephen Federowicz, Joshua A Lerman, Ali Ebrahim, Bernhard O Palsson, and Nathan E Lewis. Bigg models: A platform for integrating, standardizing and sharing genome-scale models. Nucleic acids research, 44(D1):D515–D522, 2015.

10. Zachary A King, Andreas Dräger, Ali Ebrahim, Nikolaus Sonnenschein, Nathan E Lewis, and Bernhard O Palsson. Escher: a web application for building, sharing, and embedding data-rich visualizations of biological pathways. PLoS computational biology, 11(8): e1004321, 2015.

11. Max Franz, Christian T Lopes, Gerardo Huck, Yue Dong, Onur Sumer, and Gary D Bader. Cytoscape.js: a graph theory library for visualisation and analysis. Bioinformatics, 32(2): 309–311, 2015.

12. Michael Römer, Johannes Eichner, Andreas Dräger, Clemens Wrzodek, Finja Wrzodek, and Andreas Zell. Zbit bioinformatics toolbox: a web-platform for systems biology and expression data analysis. PloS one, 11(2):e0149263, 2016.

13. Alba Lopez, Mátyás Fodor, Anna Sirunian, Ali Kaafarani, Christian Lieven, Nikolaus Sonnenschein, Tala Azrak, and Moritz E. Beber. Dd-decaf/caffeine: Version 1. https://doi.org/10.5281/zenodo.2616028, March 2019.

14. Webassembly. https://webassembly.org.

15. Benjamin J Bornstein, Sarah M Keating, Akiya Jouraku, and Michael Hucka. Libsbml: an api library for sbml. Bioinformatics, 24(6):880–881, 2008.

16. Nicolas L. Novére, Nicolas Rodriguez, Finja Wrzodek, Florian Mittag, Sebastian Fröhlich, Michael Hucka, Alex Thomas, Bernhard Ø. Palsson, Nathan E. Lewis, Andreas Dräger, Chris J. Myers, Leandro Watanabe, Ibrahim Y. Vazirabad, Victor Kofia, Harold F. Gómez, Alexander Diamantikos, Eugen Netz, Jakob Matthes, Johannes Eichner, Roland Keller, Jan Rudolph, and Clemens Wrzodek. JSBML 1.0: providing a smorgasbord of options to encode systems biology models. Bioinformatics, 31(20):3383–3386, 06 2015. ISSN 1367-4803. doi: 10.1093/bioinformatics/btv341.

17. Andreas Dräger, Nicolas Rodriguez, Marine Dumousseau, Alexander Dörr, Clemens Wrzodek, Nicolas L. Novère, Andreas Zell, and Michael Hucka. JSBML: a flexible Java library for working with SBML. Bioinformatics, 27(15):2167–2168, 06 2011. ISSN 1367-4803. doi: 10.1093/bioinformatics/btr361.

18. Alon Zakai. Emscripten: an llvm-to-javascript compiler. In Proceedings of the ACM international conference companion on Object oriented programming systems languages and applications companion, pages 301–312. ACM, 2011.

19. Andreas Rossberg, Ben L. Titzer, Andreas Haas, Derek L. Schuff, Dan Gohman, Luke Wagner, Alon Zakai, J. F. Bastien, and Michael Holman. Bringing the web up to speed with webassembly. Commun. ACM, 61(12):107–115, November 2018. ISSN 0001-0782. doi: 10.1145/3282510.

20. Webassembly garbage collection. https://github.com/WebAssembly/design/issues/1079.

21. Michael B Elowitz and Stanislas Leibler. A synthetic oscillatory network of transcriptional regulators. Nature, 403(6767):335, 2000.

22. Roland Bulirsch and Josef Stoer. Numerical treatment of ordinary differential equations by extrapolation methods. Numerische Mathematik, 8(1):1–13, 1966.

23. Gerhard Wanner and Ernst Hairer. Solving ordinary differential equations II. Springer Berlin Heidelberg, 1996.

24. Colin Smith. odex-js. https://github.com/littleredcomputer/odex-js.

25. Michael A Gibson and Jehoshua Bruck. Efficient exact stochastic simulation of chemical systems with many species and many channels. The journal of physical chemistry A, 104 (9):1876–1889, 2000.

26. Endre T Somogyi, Jean-Marie Bouteiller, James A Glazier, Matthias König, J Kyle Medley, Maciej H Swat, and Herbert M Sauro. libroadrunner: a high performance sbml simulation and analysis library. Bioinformatics, 31(20):3315–3321, 2015.

